# Fetal origin of sex-bias brain aging

**DOI:** 10.1101/2022.02.02.478867

**Authors:** Maliha Islam, Monica Strawn, Susanta K. Behura

**Author notes:** Correspondence: Susanta K. Behura.

## Abstract

DNA methylation plays crucial roles during fetal development as well as aging. Whether the aging of the brain is programmed at the fetal stage remains untested. To test this hypothesis, mouse epigenetic clock (epiclock) was profiled in fetal (gestation day 15), postnatal (day 5), and aging (week 70) brain of male and female C57BL/6J inbred mice. Data analysis showed that on week 70 the female brain was epigenetically younger than the male brain. Predictive modeling by neural network identified specific methylations in the brain at the developing stages that were predictive of epigenetic state of the brain during aging. Transcriptomic analysis showed coordinated changes in expression of epiclock genes in the fetal brain relative to placenta. Whole-genome bisulfite sequencing identified sites that were methylated both in the placenta and fetal brain in a sex-specific manner. Epiclock genes and genes associated with specific signaling pathways, primarily the gonadotropin-releasing hormone receptor (GnRHR) pathway, were associated with these sex-bias methylations in the placenta as well as fetal brain. Transcriptional crosstalk among the epiclock and GnRHR pathway genes was evident in the placenta that was maintained in the brain during development as well as aging. Collectively, these findings suggest that sex differences in the aging of brain are of fetal origin and epigenetically linked to the placenta.

## Introduction

The Dilman theory of developmental aging (DevAge) (1) suggests that mechanisms that regulate developmental processes early in life have commonalities with the processes that regulate aging. Genomics and systems-biology studies have supported the DevAge theory and have further revealed that the developmental aging link is evolutionarily conserved between mice and human (2, 3). The connection of aging with reproduction has been suggested in multiple studies (4–6). Gestation length (7), placenta development (8), and fertility (9) influences on aging and longevity. Furthermore, sex also influences differential aging of organs (10). Different organs show differential gene expression changes during aging that are sex-specific (11). In humans, the female brain remains persistently younger than the male brain (12, 13).

DNA methylation is a major factor of epigenetic regulation of aging (14–17). Methylation, the majority of which occur in CpGs (5’-Cytosine-phosphate-Guanine-3’) sites, changes in correlation with age. They are commonly referred to as epigenetic clock (epiclock) and serve as reliable predictors of the biological age of organs. Several studies have profiled methylation changes in the brain during aging (17–19) and in the developing brain at fetal stages and found that aberrant methylation in the fetal brain can influence the risk of brain diseases later in life (20, 21). Commonalities in morphometric changes have also been observed in the brain during development and aging (22). These studies have collectively suggested that processes that regulate the development and aging of the brain are not mutually exclusive. Mice models have provided further insights into epigenetic regulation of brain during development and aging (23–26), including influence of sex on epigenetic control of brain development and function (27–29).

While the role of placenta in fetal programming is well documented (30–34), recent studies show that the placenta plays adaptive roles to safeguard fetal brain development from adverse conditions during pregnancy (35). Our earlier study showed a coordinated changes of specific receptor and ligand genes between the placenta and fetal brain, suggesting their roles in the regulation of the brain-placental axis in mice (36). In addition, we also found evidences for differential expression of specific aging genes in the male and female fetal brain in response to ablation of uterine *Foxa2*, an essential gene required for pregnancy establishment (37). We also showed that specific genes are regulated in a sex-bias manner during fetal brain development in pigs (38). Furthermore, studies by others have shown that epigenetic and metabolic changes in the fetal brain can impact overall health later in life (34, 35) particularly in the context of declining metabolism in the brain (36, 37). However, the epigenetic connection between development and aging of brain remains poorly studied. The present study aims to test if development and aging of brain have epigenetic links and determine if such links are influenced by fetal sex and placenta.

## Materials and Methods

### Ethics statement

All animal procedures were approved by the Institutional Animal Care and Use Committee of the University of Missouri and were conducted according to the National Institute of Health Guide for the Care and Use of Laboratory Animals.

### Animals

Adult C57BL/6J mice were mated. The day vaginal plug was observed that was considered as gestation day (GD) 1. On GD15, dams were euthanized in CO2 followed by cervical dislocation. We selected GD15 because the placenta is fully developed, and the brain-placental crosstalk was confirmed at this time point (36). The placenta was carefully separated from the metrial gland and decidua, and the fetal brain was collected using curved forceps. All the samples were washed in PBS and snap-frozen in liquid nitrogen. Sex was determined by PCR (39). A total of 12 fetal brains and placenta were collected (3 replicates x 2 tissue types x 2 sexes). We further collected three replicates of male and female brains from postnatal day 5 (PND5) and 70 weeks old mice (WK70).

### Epigenetic clock profiling

Mouse multi-tissue epigenetic clock was profiled with DNA of GD15, PND5 and WK70 male and female brains using the Zymo (ZYMO RESEARCH, Irvine, CA 92614, U.S.A) DNAge® estimation service, which is based the Horvath pan-tissue clock using elastic net regression (40). Briefly, DNA from frozen brain samples was purified using the Quick-DNATM Miniprep Plus kit (Cat. No. D4068). Bisulfite conversion was performed using the EZ DNA Methylation-LightningTM Kit (Cat. No. D5030), followed by enrichment for target loci and sequencing on an Illumina® HiSeq instrument. Sequence reads were identified using Illumina base calling software and aligned to the mouse reference genome (GRCm38) using *Bismark* (41) which was also used for methylation calling. Methylation level was estimated as the proportion of reads mapped to each cytosine relative to the total number mapped reads to the site. The methylation data of each brain sample was then compared with cortex aging data of mouse by Zymo’s DNAge® predictor tool as described earlier (42).

### Whole-genome bisulfite sequencing (WGBS)

Genomic DNA was extracted from GD15 fetal brain and placenta of both sexes using G*entra Puregene Tissue Kit* (catalog #158667, Qiagen) as per the manufacturer’s instruction. Equimolar amounts of DNA from the three biological replicates were pooled together for each sample that used for methylation analysis as described earlier (43). After bisulfite conversion of DNA, sequencing libraries were generated using the NEBNext® Ultra™ II DNA Library Prep Kit (New England Biolabs, MA). Library preparation and sequencing were performed at the University of Missouri Genomics Technology Core. The libraries were sequenced to 20x genome coverage (150 bases sequence reads) using NovaSeq 6000.

### WGBS data analysis

The raw data were subjected to quality control by *TrimGalore* followed by mapping to the reference genome (GRCm38) by *Bismark* (41). Briefly, the reference GRCm38 genome sequences were bisulfite converted and indexed using the ‘*bismark_genome_preparation*’ function before mapping. The read alignment was then performed by *bowtie2* aligner implemented within *Bismark* (44). The methylation sites were extracted using the*bismark_methylation_extractor* to generate methylation coverage of each sample in a genome-wide manner. The coverage files contained chromosome name, start position, end position, methylation percentage, and number of methylated, and unmethylated reads for each CpG site. We only focused on CpG sites as they represent the majority of methylations in mammalian genomes (45). The sites that had low coverage (read counts < 8) were excluded from further analysis (46). The count data was converted to beta-values of methylation as described earlier (47). The raw and processed data are available in GEO accession# GSE157553.

### Genomic distribution of methylation sites

*BEDTools* (48) was used to identify methylations in different genomic locations based on genome annotation (GRCm38) obtained from UCSC Genome Browser. In addition, *awk* and *grep* were used to filter and count methylation sites in different annotation features.

### RNA-seq

RNA-seq was performed to profile gene expression of male and female brains collected on PND5 and WK70. The RNA-seq data of brain and placenta from male and female fetuses (GD15) was compared with RNA-seq data of PND5 and WK70 brain. The GD15 RNA-seq data was generated earlier by us (37) which are also publicly available in the Gene Expression Omnibus database (accession # GSE157555). Total RNA was isolated from PND5 and WK70 brain samples using TRIzol (Catalog 15596026, Thermo Fisher) based protocol. The RNA quality was checked using a Fragment Analyzer (Advanced Analytical Technologies). The RNA concentration was determined using a Qubit 2.0 Fluorometer (Life Technologies). Libraries were prepared from total RNA using the Illumina TruSeq Stranded Total RNA with Ribo-Zero Gold Library Prep Kit at the University of Missouri Genomics Technology Core. Each library was sequenced (∼30 million paired-end reads, 150 bases read length) using Illumina NovaSeq 6000 sequencer.

### RNA-seq data analysis

RNA-seq data analysis was performed as described in our earlier studies (37, 38). Briefly, the quality of raw sequences was checked with FastQC followed by trimming the adaptors from the sequence reads by *cutadapt*. The *fqtrim* tool was used to perform base quality trimming (Phred score >30) by sliding window scan (6 nucleotides). The quality reads were then mapped to the mouse reference genome GRCm38 using *Hisat2* aligner (49). Read counting from the alignment data was performed by *FeatureCounts* (50).

### Mutual information network analysis

The information theory (MI) approach (51) was used to infer gene expression networks. MI is a measure of the information content between two variables: a numerical value ranging from 0 to 1 depending on, intuitively, how much knowing one variable would predict the variability of the other. We calculated MIs in a pair-wise manner, both at gene and sample levels, to generate a weighted adjacency matrix by the Maximum Relevance Minimum Redundancy (MRMR) method (52). Then mutual information network analysis was performed using *minet* (53) followed by network visualization by *GGally*.

### Statistical analysis

All statistical tests were performed in *R*. Chi-Square analysis was performed to infer the significance of 2×2 contingency tests. Enrichment analysis was performed by Fisher exact tests followed by multiple corrections of *p*-values by False Discovery Rate (FDR). Comparison of variation between methylation and expression was based on beta-values of methylation (47) and fragment per kilobase per million (FPKM) values of expression (54). Distances calculation of methylation and expression variation was based on the *Euclidean* method. Hierarchical clustering was performed from the distance measures by *ward*.*D2* as the method of agglomeration. Comparison between clusters (*tanglegram* analysis) was performed using the package *dendextend*. The canonical correlation analysis was performed by *cancor* function of *CCA* package. In this analysis, expression of epiclock genes in the GD15 placenta was treated as the independent variable (x-values), and the fetal brain was treated as the dependent variable (y-values). Correlation coefficients (coefficients of x and y values) were multiplied with the respective data matrices of fetal brain and placenta to generate covariates for comparison. The *neuralnet* package was used to train neural network (NN) models by backpropagation using the weight backtracking method (55). NN consists of three layers: input-layer that takes training data for model to learn, hidden-layer uses backpropagation to optimize the weights of the input variables to develop the model, output-layer uses the trained model to predicts the variable from a test data. The methylation data of the aging brain was used as predicted values, and those of fetal and postnatal brains were used as predictor values in NN modeling. The trained model was used to separately predict the methylation level in male and female aging brains. Finally, the observed and model-predicted methylation values of the aging brain in both sexes were compared to create confusion matrix using *caret* package to evaluate model accuracy.

## Results

### Epigenetic clock profiles of male and female brain

DNA methylation of mouse epiclock was profiled in GD15, PND5, and WK70 male and female brains (**Supplementary Table 1**). The methylation data were used in epigenetic age (also called DNA age) analysis (56) which showed that the female brain was biologically younger (29.6 weeks) than the male brain (38.1 weeks) despite the same chronological age of the animals. The female brain showed a higher level of methylation than the male brain for 854 sites, whereas the male brain showed higher methylation than the female brain for 1,028 sites (**Figure 1a**). A relatively lower level of epiclock methylation was observed in the fetal and postnatal brain compared to the aging brain (**Supplementary Figure 1**). Methylation level either increased or decreased in the brain among the three life stages (fetal, postnatal, and aging) in a sex-biased manner (**Figure 1b**). No significant bias was observed in a 2×2 contingency analysis of the number of sites that either increased or decreased in methylation level of male and female brain at fetal versus postnatal stage (*P* =0.84). But a significant (*P* < 0.0001) difference was observed between postnatal and aging stage. Upon aging, a greater number of sites showed increased methylation in the brain of males than that of females (**Figure 1b**). Furthermore, specific epiclock genes were methylated in a sex-biased manner at all three life stages. A total of 245 epiclock sites showed increased methylation in the female brain than the male brain at each of the three stages (**Figure 1c**). An opposite pattern was observed with 187 sites (**Figure 1d)**. The female bias methylations showed higher median beta-values (methylation level) compared to that of male bias methylation sites (**Figure 1d**). Moreover, the female bias methylations showed higher level of variation than that of male bias methylations, and this pattern was consistent among the fetal, postnatal, and aging stages.

**Table 1.** Number of FB and MB methylations in fetal brain and placenta in different genic and intergenic features of the genome.

| **Feature** | **FB** | **MB** |
| --- | --- | --- |
| Exon | 662 | 730 |
| Intron | 7594 | 8141 |
| UTRs | 298 | 331 |
| Promoter | 3 | 4 |
| CpG island | 1 | 3 |
| CpG shore | 165 | 175 |
| Refseq Functional Element | 14 | 20 |

**Figure 1.**
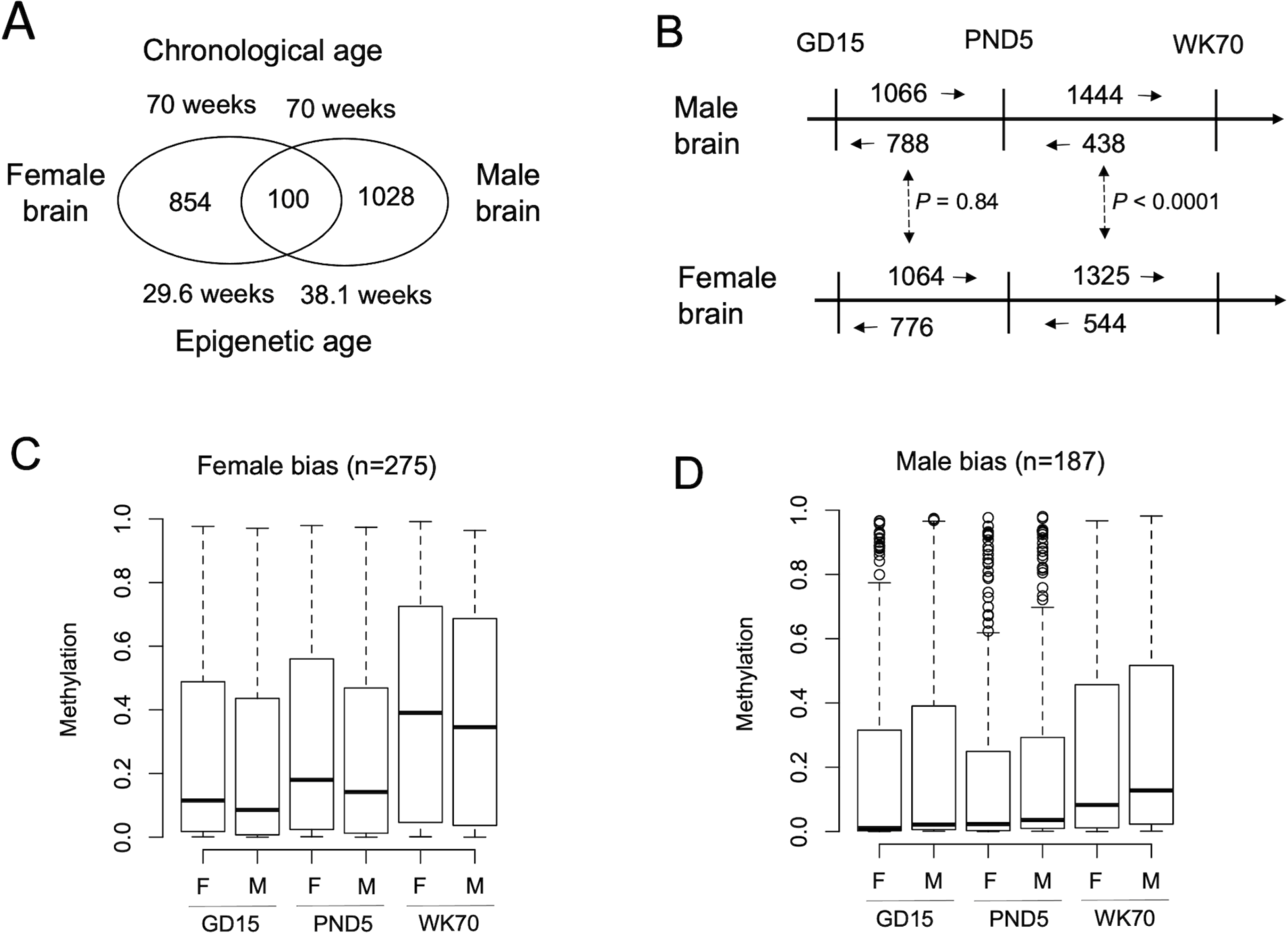
Epigenetic clock analysis of aged and developing brain in male and female mice. **A**. Epigenetic age, based on the methylation status of the epigenetic clock, of the male and female brain of 70 weeks old mice. The Venn diagram shows the number of epiclock sites with higher methylation in females than males, or *vice versa*. The intersection shows the number of sites that had the same level of methylation in the brain in both sexes. **B**. Comparison of methylation of epiclock in the male and female brain at fetal stage (GD15) and postnatal stage (PND5) with the aged stage (WK70). The number and direction of arrows show a higher level of methylation of epiclock sites between stages (arrowhead shows the stage where methylation is higher). 2×2 contingency test shows no significant bias in the number of CpGs with increase or decrease methylation during brain development (GD15 vs. PND5) but significant bias between PND5 and WK70. **C**. Number of epiclock sites showing consistently either female bias or male bias methylation in the brain among all three life stages (fetal, postnatal, and aging). **D**. Boxplot shows the female-bias epiclock methylation in the brain among the three life stages. The median methylation is shown by horizontal lines inside the boxes. **E**. Boxplot shows the male-bias epiclock methylation among the three life stages. The median methylation is also shown by horizontal lines inside the boxes.

### Developmental methylations as predictors of aging brain

We employed NN modeling to test if methylation changes of epiclock genes in the brain at developmental stages (fetal and postnatal) are predictors of methylation of the aging brain (**Figure 2**). The epiclock methylation of the brain at fetal and postnatal stages were used to train NN model to learn the corresponding changes in methylation of the aging brain separately in males and females. The model converged after 778 iterative steps (number of optimizations at an error rate of less than 5%) with an average error rate of 1.69 in the female brain (**Figure 2a**). In the male brain, the model converged with only 175 iterations with an average error of 1.15 (**Figure 2b**). This agrees with our observation that variation in epiclock methylation is relatively higher in the female brain than the male brain. The confusion matrix showing the number of true and false predictions relative to the observed data are shown in **Figure 2c and d**. It shows that the model identified 113 methylations in the brain that occurred in the developmental stages (fetal and postnatal stages) that were predictive of epiclock changes in the brain upon aging (**Supplementary Table 2)**.

**Figure 2.**
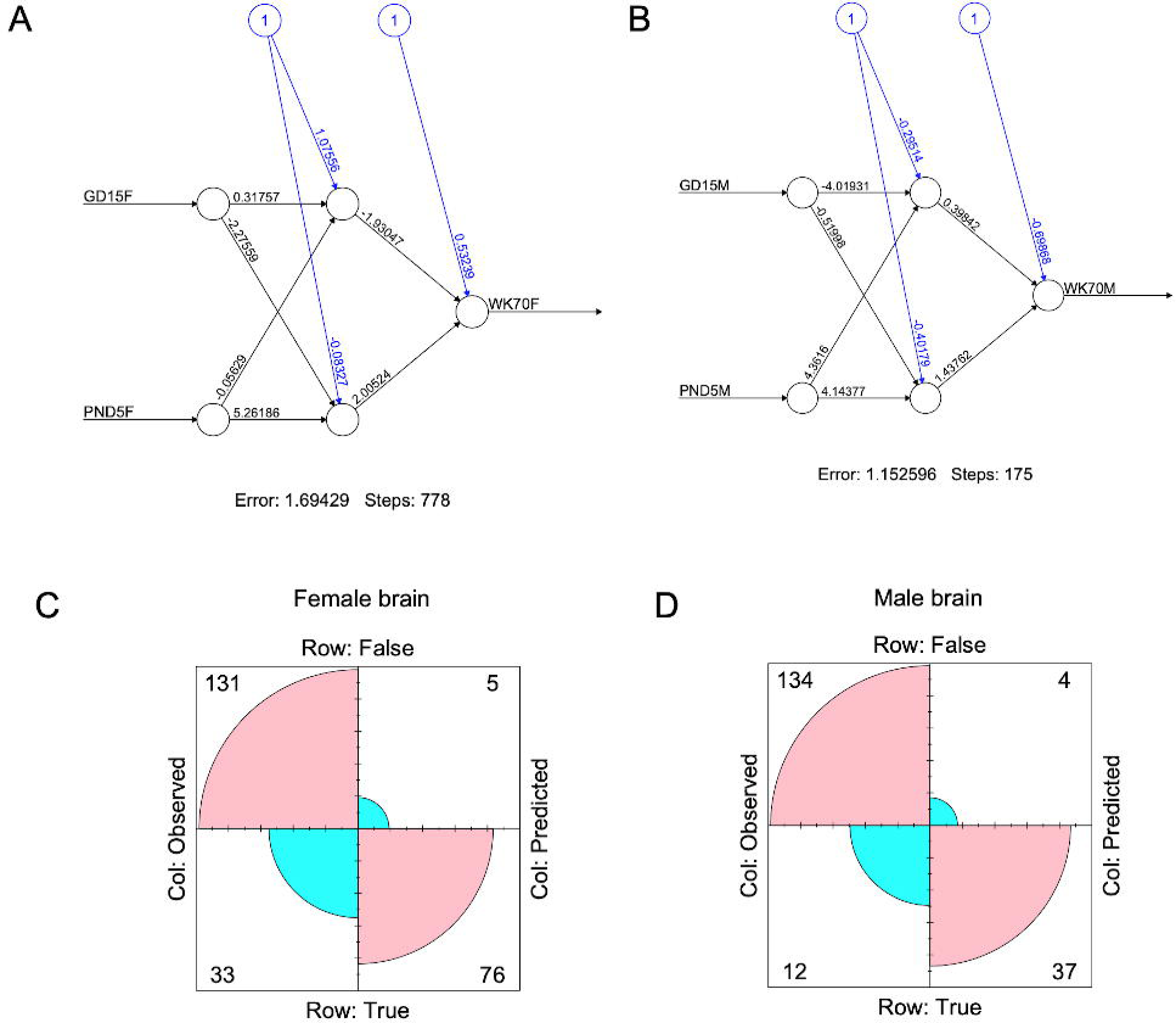
NN modeling of the epigenetic clock variation. **A**. Epigenetic clock with female-bias methylation data was used to train NN model. **B**. The epigenetic clock with male-bias methylation data was used to train NN model. The trained models were then used for predictions of methylation of aging brain. The confusion matrix shows the number of true and false predictions relative to the observed methylation data (**C** and **D)**.

### Expression changes of epiclock genes in brain

RNA-seq was performed to profile gene expression of the same brain samples used for methylation analysis. Methylation changes of the predictive markers identified from *neuralnet* analysis (listed in **Supplementary Table 2**) were compared with expression level of the cognate genes in the brain at the fetal, postnatal, and aging stages. The tanglegram in **Figure 3a** showed that methylation changes were more similar between fetal and postnatal brain than aging brain. But expression changes were more similar between postnatal and aging brain than the fetal brain. This pattern of yin and yang between methylation and expression changes in the brain suggested that low methylation was associated with higher expression in the fetal brain, whereas increase in methylation was linked to lower expression of genes upon aging (**Figure 3**b and c).

**Figure 3.**
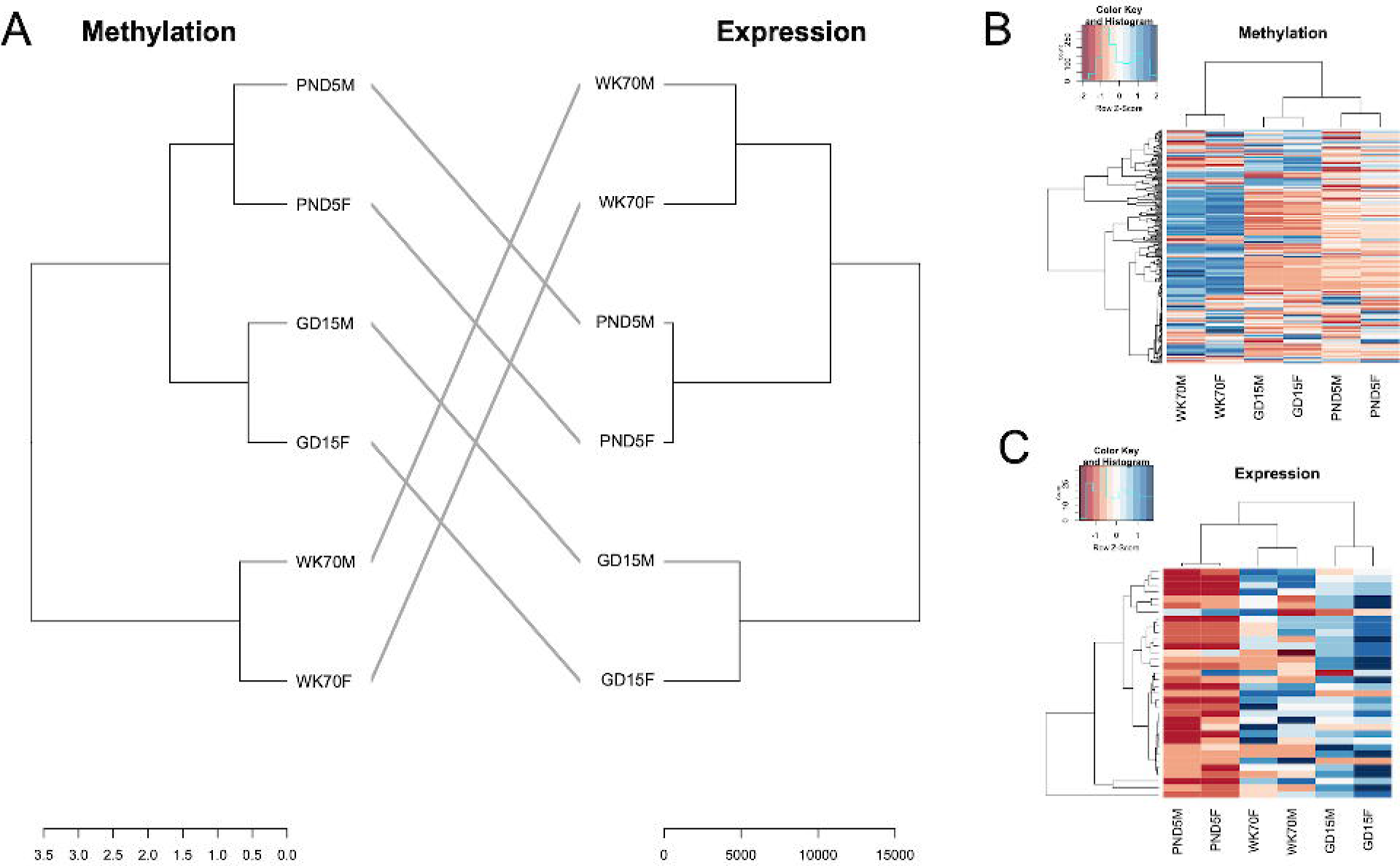
Comparison of methylation and expression variation of epiclock prediction maker genes. **A**. Tanglegram shows comparison of methylation and expression patterns of the epiclock predictive genes (from NN models) among the fetal, postnatal, and aging brains of both sexes. **B**. Heatmap of methylation variation of epiclock marker genes. **C**. Heatmap of expression variation of epiclock marker genes.

Next, we compared expression changes of the epiclock genes in the brain relative to placenta (**Figure 4a**). The hierarchical clustering analysis showed that epiclock genes were expressed in fetal brain in a more similar manner with placenta than the postnatal and aging brain. However, genes unrelated to epiclock did not show this pattern (**Figure 4b**). In non-epiclock genes, expression of fetal brain grouped together with that of postnatal and aging brain, and gene expression of placenta was distinct. This suggested that epigenetic clock is coordinately expressed between placenta and fetal brain. To further test this idea, canonical correlation analysis (CCA) (57) was performed. CCA showed that covariates of expression (which is expression multiplied with the correlation coefficient) of epiclock genes in the fetal brain was positively correlated with that of placenta (**Supplementary Figure 2**). But this relationship was absent in the expression of non-epiclock genes. This supported our idea that epigenetic clock is active during pregnancy and is likely associated with the regulation of the brain-placental axis (36, 58).

**Figure 4.**
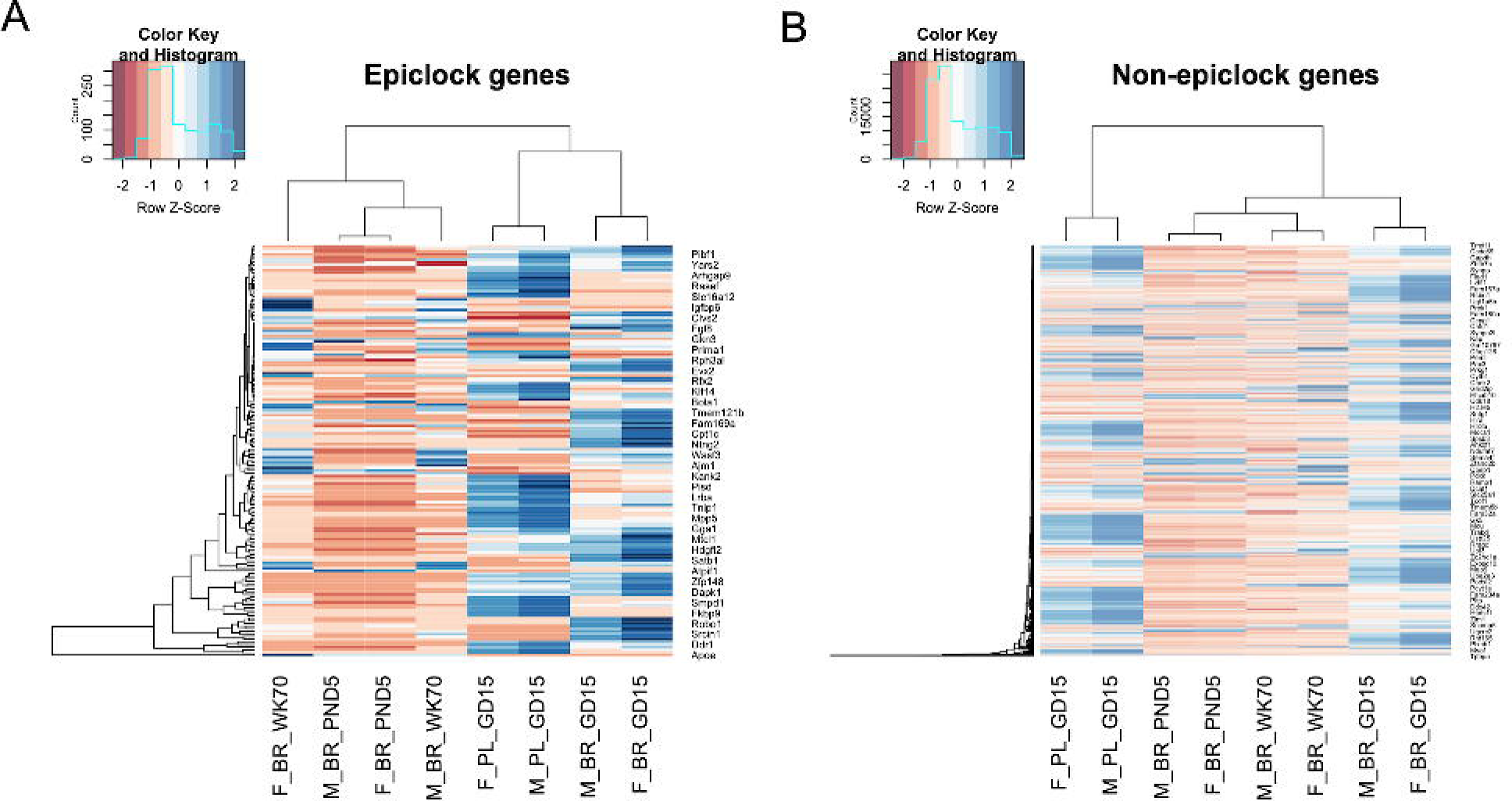
Comparison of placenta and brain gene expression. **A**. Heatmap of expression of all epiclock genes in placenta along with the fetal brain, postnatal brain, and aging brain. **B**. Heatmap of expression of non-epiclock genes in placenta and fetal brain, postnatal brain, and aging brain. In both plots, the scales on the top left show the color codes for methylation levels.

### Placenta shows sex-bias methylation in concert with fetal brain

Whole-genome bisulfite sequencing (WGBS) was performed to further analyze global methylation of placenta in a comparative manner with that of fetal brain in both sexes. WGBS (data available in GEO, accession# GSE157553) identified 15,269 and 17,028 CpGs that were methylated in female-bias (FB) and male-bias (MB) manner respectively in both fetal brain and placenta (**Figure 5**a and b). The genomic position and methylation level of these sites are provided in **Supplementary Table 3**. A significant (*P=*0.0003) bias was observed in the location of these methylations in genic versus intergenic regions in the genome (**Figure 5**c). While some genes harbored either FB or MB methylations, genes were also identified that harbored both FB and MB methylations (**Figure 5**d).

**Figure 5.**
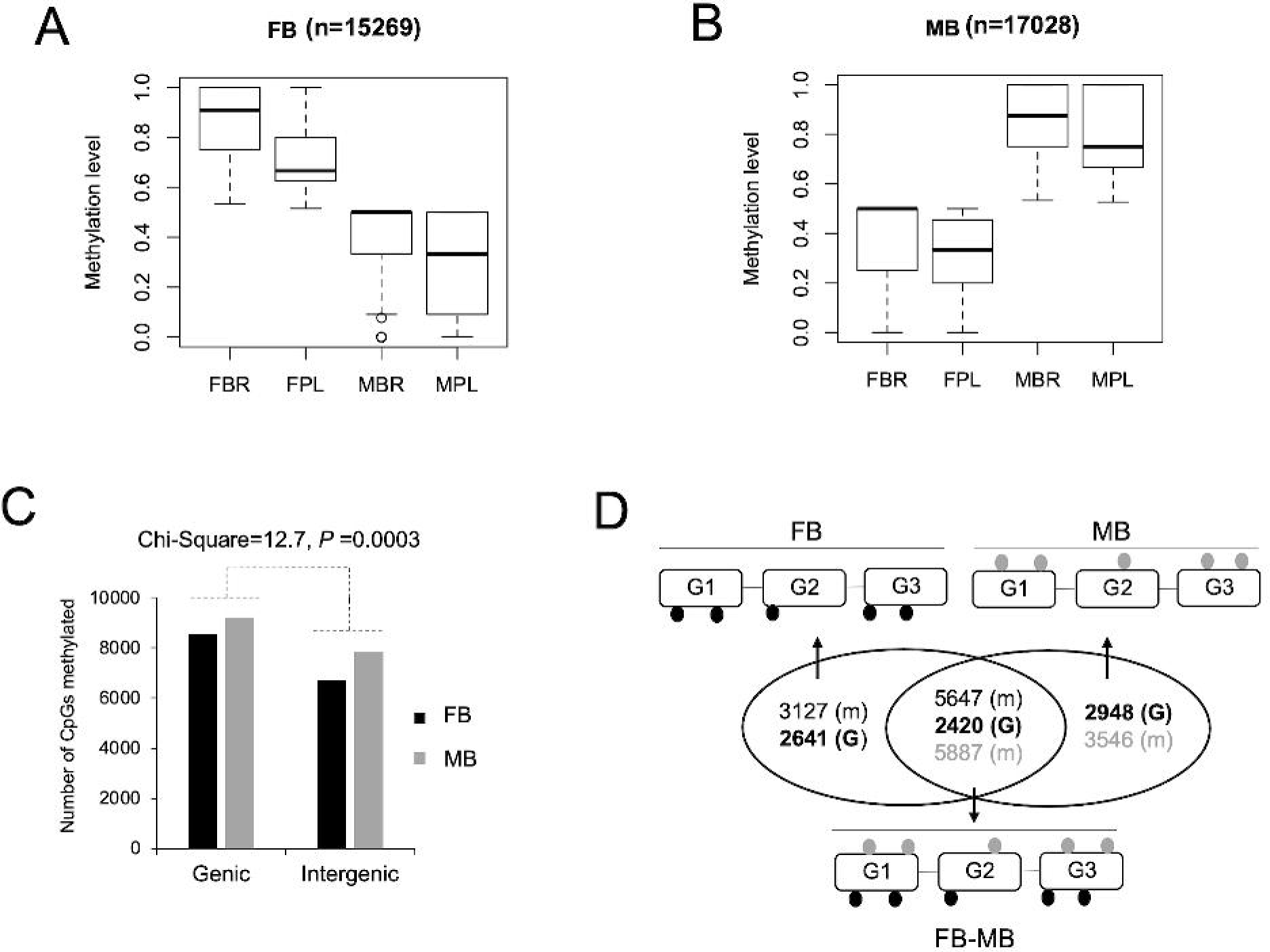
Common sex-bias methylation in the placenta and fetal brain. **A**. Boxplot showing that specific sites are methylated in a female-bias (FB) manner in both placenta and fetal brain. Y-axis shows beta-values of methylation. X-axis shows brain (BR) and placenta (PL) of both sexes (M= male and F=female). **B**. Boxplot showing that specific sites are methylated in a male-bias (MB) manner in both placenta and fetal brain. Y-axis shows beta-values of methylation. X-axis shows brain (BR) and placenta (PL) of both sexes (M= male and F=female). **C**. Barplot showing the number of methylations that were either female-bias or male-bias methylations within genes or in intergenic regions. The 2×2 contingency test *p*-value of association (sex vs. location) is shown. **D**. Venn diagram showing the number of genes with either male-bias or female-bias methylation or both male-as well as female-bias methylations. The cartoons show the location of male and female bias methylations in those genes for illustrative purpose.

The sex-bias methylations in gene bodies were predominantly found in introns. They were roughly 11-fold more in number than those in exons (**Table 1**). The frequency of sex-bias methylations was low in the untranslated regions (UTRs). While we identified 630 FB and 587 MB methylations within *cis*-regulatory regions, we rarely found these methylations in the promoters and Refseq functional elements (59). Sex-bias methylations were more frequent in CpG shores than CpG islands (**Table 1**).

Additionally, sex-bias methylations were found within repeat elements, the majority of which were associated with different retrotransposons both within genic and intergenic regions. While, in general, methylations occurred at a higher level in repeat sequences than non-repeat sequences in the genome, the number of sex-bias methylations were significantly different genic versus intergenic repeats (**Figure 6**) indicating a role of heterochromatin methylation in sex-bias gene regulation between fetal brain and placenta (28, 60–62).

**Figure 6.**
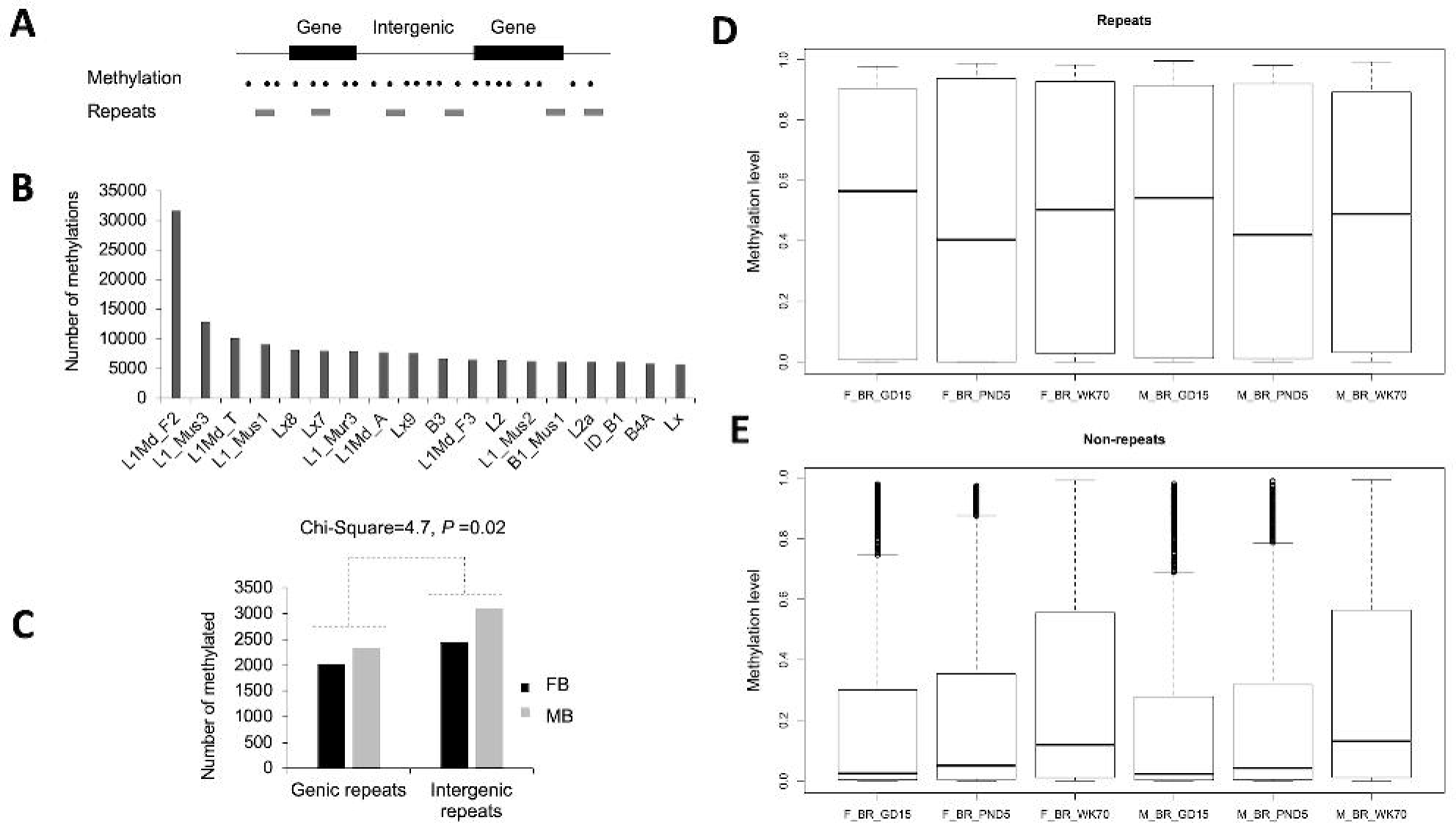
Methylation of repeat elements in the placenta and fetal brain. **A**. A schematic representation of methylation of repeat elements either in genic or intergenic regions. **B**. Barplot showing the number of all methylations (sex-bias or not) in different repeat elements. Top-most abundant repeats (based on number of methylation) are shown. **C**. Number of sex bias methylations observed in both placenta and fetal brain are differentially associated (significance level shown) with repeat elements within genic and intergenic regions. **D**. Comparison of methylation level of epiclock sites within repeat elements among fetal, postnatal, and aged brain. **E**. Comparison of the methylation level of epiclock sites found within non-repeat elements among fetal, postnatal, and aged brain.

Functional annotation of genes harboring the sex-bias methylations was performed. The analysis showed significant enrichment of specific pathways (Fisher exact test, *P* < 9.05) most of which were related to signaling (**Supplementary Table 4)**. Genes with FB methylations were enriched with T cell activation, Huntington disease, FGF and Cadherin signaling pathways. Genes with MB methylations were enriched with pathways of axon guidance mediated by netrin, histamine H1 receptor mediated signaling, thyrotropin-releasing hormone receptor signaling, and heterotrimeric G-protein signaling. Also, specific pathways were commonly enriched by genes harboring both FB and MB methylations (see **Supplementary Table 4)**.

### Crosstalk between epiclock and signaling genes in fetal brain is maintained in aging brain

We identified specific epiclock gens and signaling pathway genes (**Supplementary Table 5**) that were methylated in both placenta and fetal brain in a coordinated manner. Principal component analysis showed two distinct groups of methylations in these epiclock and signaling genes identified in both placenta and fetal brain (**Figure 7a)**. Hierarchical cluster analysis showed specific epiclock genes (*Khdrbs2, Brinp2, Clvs2, Sfi1, Gas7, Rhot1, Bcas3, Ankrd24, Sox30, Trim7, Wnt3a*, and *Lrrc75a*) and gonadotropin-releasing hormone receptor (GnRHR) pathway genes (*Pbx1, Atf3, Rap1b, Fos, Grb2, Zeb1, Smad4, Cdc42*, and *Raf1*) were coordinately methylated in both placenta and fetal brain in a sex-dependent manner (**Figure 7**b and c). The sex-bias methylations of these genes are listed in **Supplementary Table 5**. Mutual information network analysis (51) of expression changes of these genes further showed that they interacted differentially in males compared to females (**Figure 8**a and b). Furthermore, the analysis showed that the expression of these genes in the placenta was mutually informative in the brain at fetal as well as postnatal and aging stages (**Figure 8**c and d). This suggested that crosstalk among the epiclock genes and GnRHR pathway genes in the placenta is epigenetically maintained in the brain throughout life, but in a sex-bias manner.

**Figure 7.**
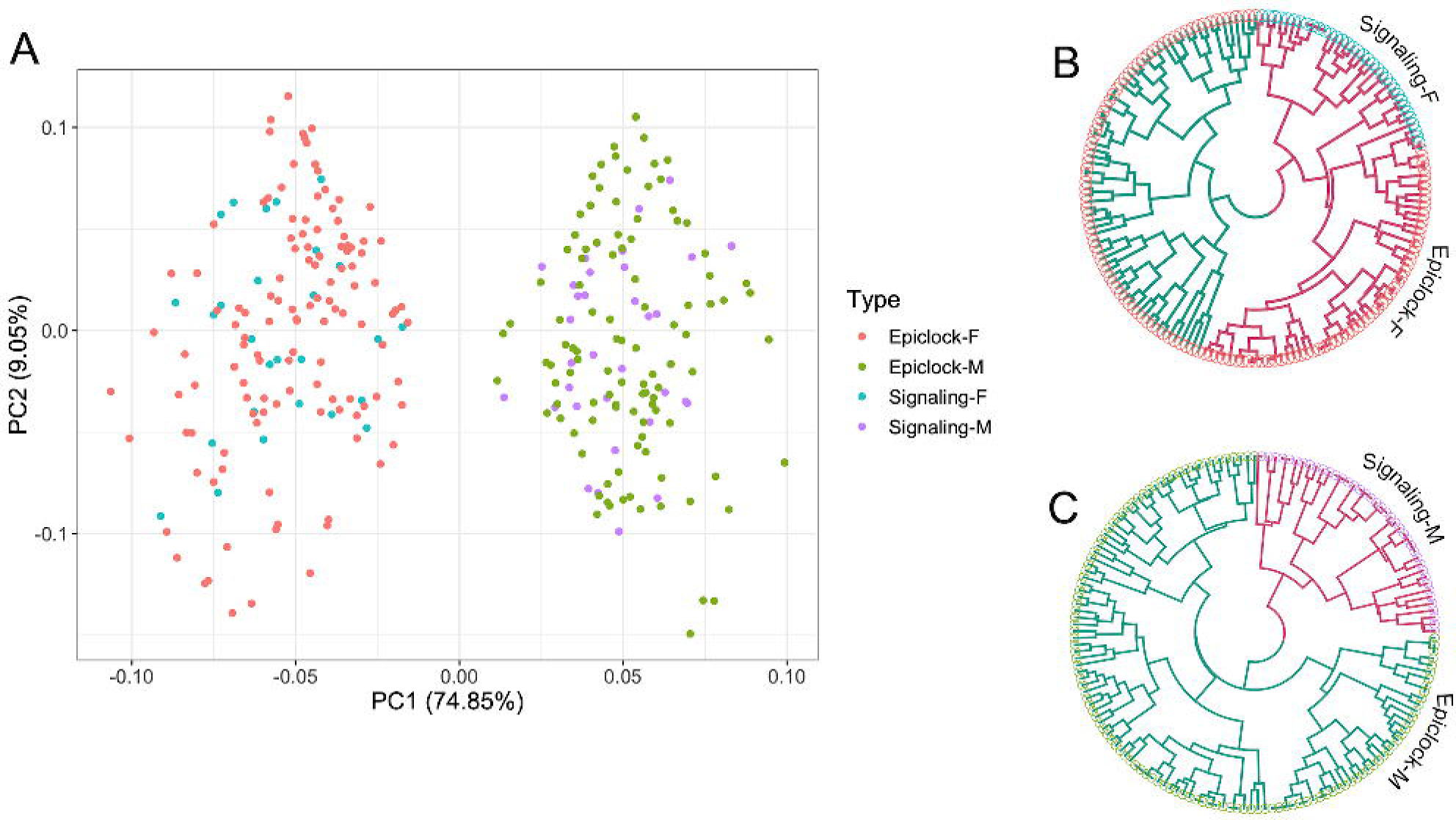
Sex-bias methylation changes among epiclock and signaling genes in the brain. **A**. Principal component analysis of male (M) and female (F) bias changes in methylation of epiclock and signaling genes in the brain among the fetal, postnatal, and aging stages. The color codes are according to the association (types) of the methylations with genes and sex listed in the legend right to the plot. **B**. Circular dendrogram based on hierarchical cluster analysis of female-bias methylations. The leaf color in the dendrogram matches to color code description in **A**. The branch color of the two clusters is shown in red and green. **C**. Circular dendrogram based on hierarchical cluster analysis of male-bias methylations. The leaf color in the dendrogram matches to color code description in **A**. The branch color of the two clusters is shown in red and green.

**Figure 8.**
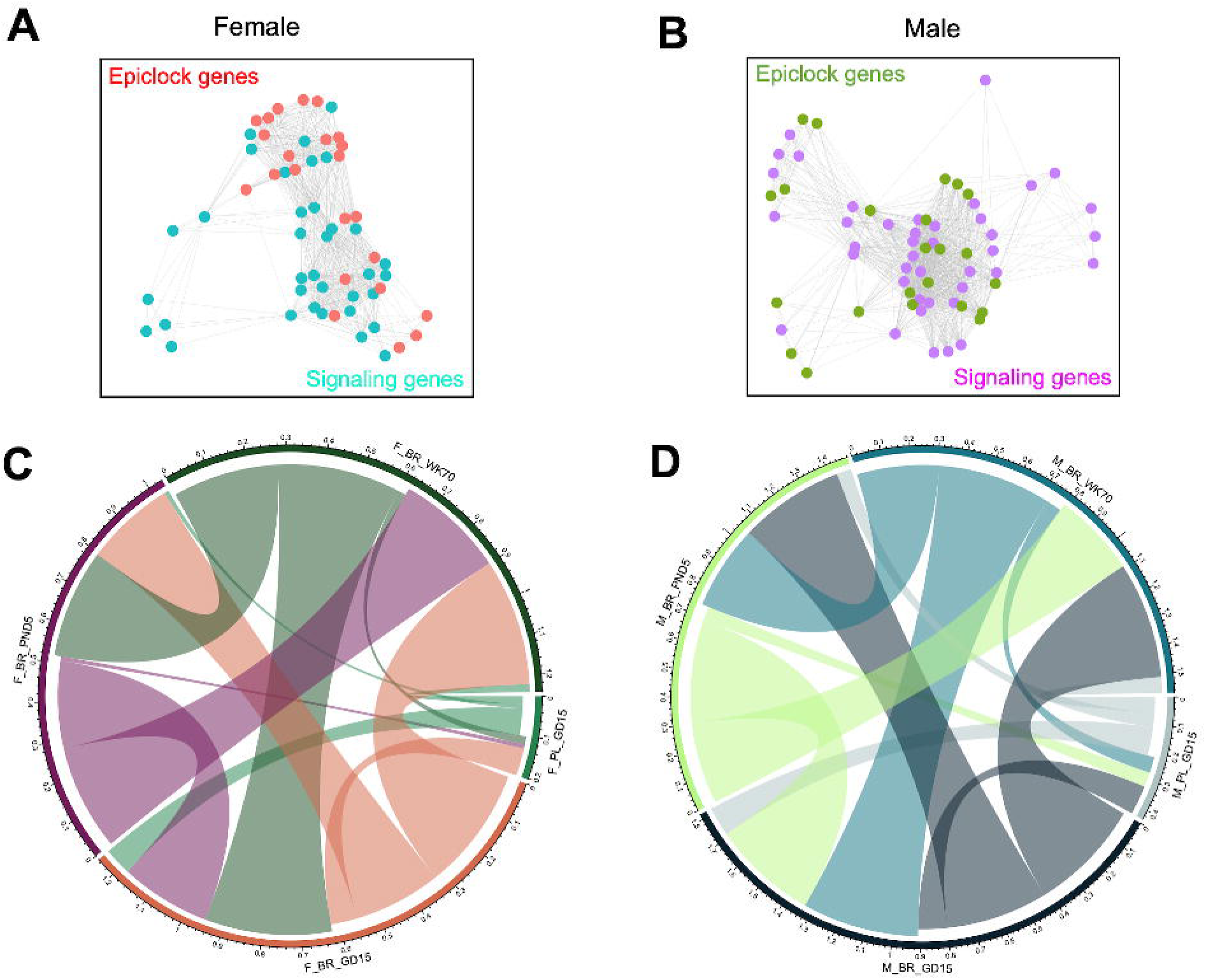
Crosstalk among epiclock and signaling genes. Expression network among epiclock and signaling genes associated with female-bias (**A**) and male-bias methylations (**B**). Color codes represent gene types shown in the plots. **C**. Circos plot showing placental links with brain in females. **D**. Circos plot showing placental links with brain in males. The arches in both C and D represent the information contents mutually shared between the placenta and brain (shown on the circumference of the plots).

## Discussion

The current study investigated the epigenetic aging of male and female brain of C57BL/6J mice. The male brain showed higher epigenetic age than that of female brain. As the chronological age was same (70 weeks), this suggested that the epigenetic aging of the brain is sexually dimorphic. Sexual dimorphisms of brain have been reported in diverse animals, including flies (63), birds (64), rodents and humans (65). Sex differences in brain aging is also known (66–70). Results from the current study in mice are consistent with earlier study in humans which showed that the female brain remains younger than the male brain in a persistent manner (12). Evolutionary studies suggest that natural selection may shape brain neoteny (youth), and such selective forces can vary within and between species (67, 71). Moreover, brain aging can vary at individual as well as population levels within species (72, 73). Such heterogeneity in brain aging have been well documented (74–76). This includes studies in mice that showed that the inbred mice strains are age differentially (77). In addition, influence of sex on aging has also been implicated (78).

DNA methylation changes in the brain throughout life (18, 79, 80). Our data showed that there are patterns how these changes in brain as mice age. We found that specific CpG were persistently methylated in a sex-bias manner in fetal, postnatal, and old age. Epigenetic clock analysis (81) identified CpGs (n=275) that were methylated higher in females than males. In contrast, relatively fewer sites (n=187) were methylated higher in males than females. The female bias methylations had a higher level of variation than male bias methylations in the brain at all stages (**Figure 1**) which supported earlier observation that persistent methylations were required in female brain to suppress genes associated with brain masculinization (82). We were further able to identify specific epiclock methylation and associated genes as early-life predictors of brain aging in females versus males.

Methylation impacted gene expression in the brain (**Figure 3**). Methylation level was lower in the brain at fetal and postnatal stages than old age. In contrast, expression level was higher in postnatal and old ages than in fetal stage. This contrasting pattern suggested that methylation likely suppressed epiclock gene expression in the brain starting after birth. An earlier study in humans has shown that the brain transcriptome undergoes a significant remodeling at the postnatal age (71). Additionally, we observed that epiclock expression of the fetal brain was more related to that of the placenta than that of the postnatal and aging brain. Genes not related to epiclock did not this pattern. Expression of those genes in the fetal brain was more related to postnatal and aging brain than the placenta. This suggested that epiclock genes are regulated between placenta and fetal brain in a manner that is distinct from all other genes active during pregnancy.

We performed whole-genome methylation analysis that showed large distinctions in DNA methylation between fetal brain and placenta. It was expected as tissue-specific variation in DNA methylation has been extensively reported from previous studies (83– 85). While different tissues accumulate methylations differentially, concordance in DNA methylation between tissues are also known (86). Studies have shown concordance in DNA methylation in mouse placenta and fetal brain in response to exposure to polychlorinated biphenyls (87) and prenatal stress (88), indicating possible epigenetic programming of fetal brain by the placenta. In the current study, we observed that both placenta and fetal brain were coordinately methylated specific CpG sites in a sex-bias manner (**Figure 5**). These methylations occurred in gene bodies more often than in intergenic regions, which may support earlier reports that gene body methylations play essential roles in placental functions (89, 90). We observed that the sex-bias methylations were higher in repeat elements including L1 retrotransposons, *Alu* repeats and short interspersed repeats (**Figure 6**) than methylation in non-repeat sequences further supporting that heterochromatin methylation may have common influence on reproduction (91) and brain development (17).

A key finding of this study was the evidence of crosstalk among epigenetic clock and GnRHR pathway genes in the placenta and fetal brain (**Supplementary Table 5**). Network analysis showed evidence for placental links to brain that is maintained in the brain at postnatal and the aging stage. GnRH plays key roles in mammalian reproduction by inducing luteinizing hormone (LH), and follicle-stimulating hormone (FSH). It also functions in DNA modification that is required to express the target genes in gonadotrope (92). Besides a role in the anterior pituitary, GnRH also plays important roles in the placenta, likely to modulate the maternal-fetal interface (93). In mice, gonadotropin production varies in a sex-dependent manner during late gestation, in which fetal androgen plays a critical regulatory function (94). Regulation of GnRH is suppressed by androgen in males, which plays roles in brain masculinization (95). Suppression of genes associated with brain masculinization is mediated by DNA methylation in the brain of females (82). Multiple studies have shown that exposure to sex hormone at the fetal stage plays a leading role in sex-specific fetal programming of cellular metabolism that poses a differential risk to metabolic diseases later in life (96, 97), including a metabolic decline of the brain (98).

In conclusion, the current study shows that epigenetic aging of brain is sex-bias, and further suggests that this bias is linked to the placenta as evident from DNA methylation of placenta relative to the brain of male versus female fetuses. The results support DevAge theory (1) that early-life developmental processes have epigenetic links to aging processes. It further highlights that aging is a biological process that originates from the fetal stage and persists throughout the life.

## Acknowledgments

The authors are thankful to Dr. Pramod Dhakal for assisting fetal brain dissections. Nathan J. Bivens, Mingyi Zhou, and Christopher Bottoms of Genomics Technology Core provided help in library preparation, sequencing, and raw data processing.

## Supplementary Figure legends

**Supplementary Figure 1**. Heatmap showing variation in methylation of epigenetic clock in the brain of males and females at GD15, PND5, and WK70. The scale on the top left shows the color codes for methylation levels.

**Supplementary Figure 2**. Scatter plots showing the relationship between covariates (of two major axes x and y calculated from the canonical correlation analysis) of methylation changes in placenta relative to the fetal brain of both sexes. The pattern for epiclock genes is shown in **A**, and for all other genes is shown in **B**.

## Supplementary Table List

**Supplementary Table 1**. Methylation level (beta-value) of mouse epigenetic clock in fetal (GD15), postnatal (PND5), and aging (WK70) brain of female and male mice. Associated genes are also shown.

**Supplementary Table 2**. List of early-life methylation marks that are predictive of brain aging in males and females.

**Supplementary Table 3**. List of CpG sites showing sex-bias methylation in both placenta and fetal brain. The methylation level in the placenta and fetal brain are shown. Genes associated with methylations are also listed.

**Supplementary Table 4**. Significant enrichment of specific pathways by genes with either male-or female-bias methylations in the placenta and fetal brain.

**Supplementary Table 5**. List of methylations in epiclock and signaling genes that show differential crosstalk in males versus females. The epiclock genes that were identified as predictive markers of brain aging and the genes associated with the GnRHR pathway are indicated.

