## Supplementary figures and images for "Fetal origin of sex-bias brain aging"

### Supplementary Figure 1

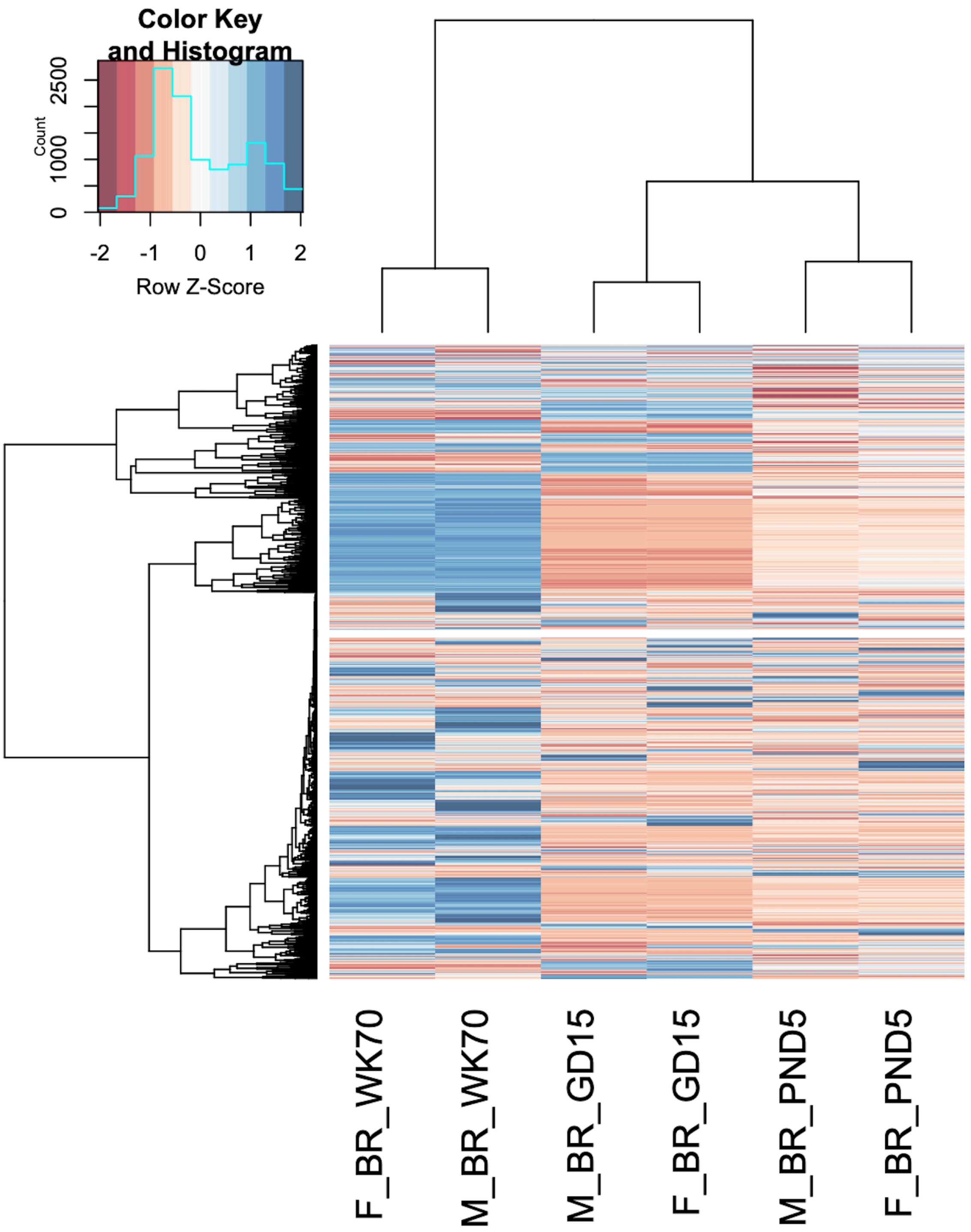

### Supplementary Figure 2

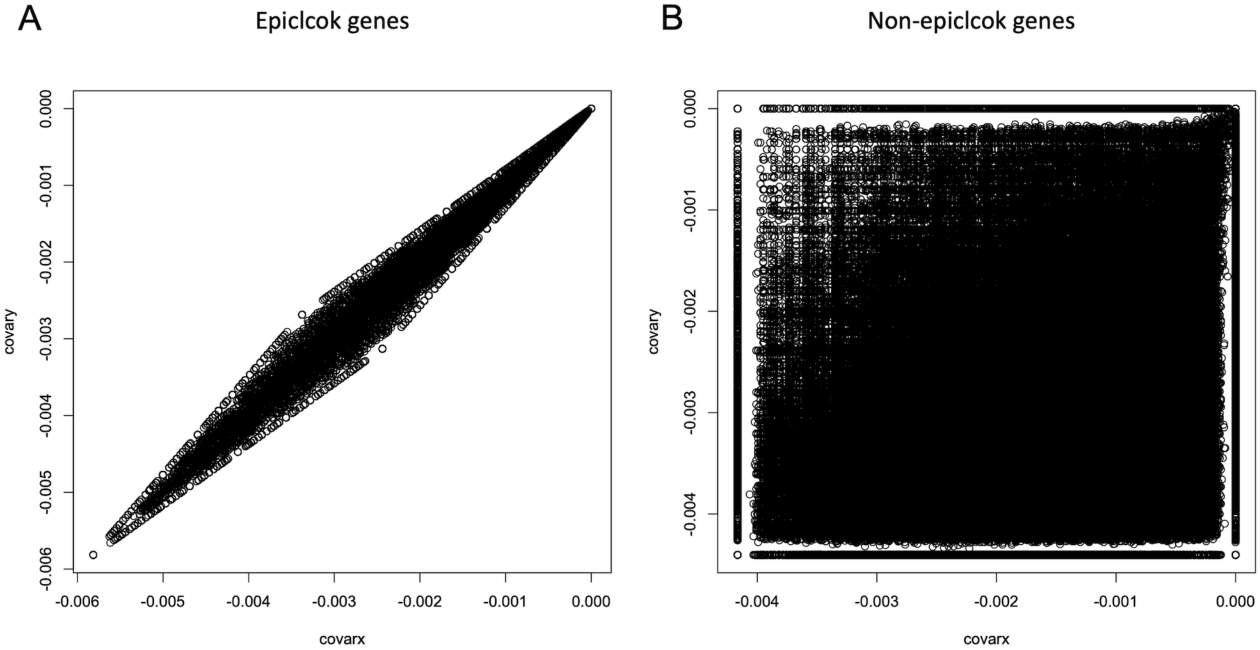
